# A cell-based evaluation of a non-essential amino acid formulation as a non-bioactive control for activation and stimulation of muscle protein synthesis using *ex vivo* human serum

**DOI:** 10.1101/713768

**Authors:** Bijal Patel, Martina Pauk, Miryam Amigo-Benavent, Alice B. Nongonierma, Richard J. Fitzgerald, Philip M. Jakeman, Brian P. Carson

**Author notes:** **Corresponding Author:** Brian P. Carson, PhD, Department of Physical Education & Sport Sciences, Faculty of Education and Health Sciences, University of Limerick, Ireland. **Conflict of interest:** The authors declare no conflict of interest.

## Abstract

**Purpose:** The purpose of this study was to compare the effect of treating skeletal muscle cells with media conditioned by postprandial *ex vivo* human serum fed with either isonitrogenous NEAA or a whey protein hydrolysate (WPH) on stimulating MPS in C2C12 skeletal muscle cells.

**Methods:** Blood was taken from six young healthy males following overnight fast (fasted) and 60 min postprandial (fed) ingestion of either WPH or NEAA (0.33 g.kg^-1^ Body Mass). C2C12 myotubes were treated with media conditioned by *ex vivo* human serum (20%) for 4 h. Activation of MPS signalling (phosphorylation of mTOR, P70S6K and 4E-BP1) were determined *in vitro* by Western Blot and subsequent *de novo* MPS were determined *in vitro* by Western Blot and surface sensing of translation technique (SUnSET) techniques, respectively.

**Results:** Media conditioned by NEAA fed serum had no effect on protein signalling or MPS compared to fasted, whereas media conditioned by WPH fed serum significantly increased mTOR, P70S6K and 4E-BP1 phosphorylation (p<0.01, p<0.05) compared to fasted serum. Furthermore, the effect of media conditioned by WPH fed serum on protein signalling and MPS was significantly increased (p<0.01, p<0.05) compared to NEAA fed serum.

**Conclusion:** In summary, media conditioned by NEAA fed serum did not result in activation of MPS. Therefore, these *in vitro* findings suggest the use of isonitrogenous NEAA acts as an effective control for comparing bioactivity of different proteins on activation of MPS. These findings also confirm that activation of MPS in C2C12 myotubes treated with media conditioned by WPH-fed serum is primarily due to circulating EAA.

## 1. Introduction

Muscle protein synthesis (MPS) is integral to the repair, growth and maintenance of skeletal muscle and sensitive to nutrient ingestion. Several studies have assessed the role of protein and amino acids in the regulation of MPS (1-4). The importance of appropriate controls in establishing the bioactivity of compounds in human MPS studies has been emphasised in recent reviews by Morton et al. (5) and Phillips (6). These meta-analyses include many studies reporting on the effects of amino acid and protein supplementation on MPS which use either a less appropriate (including carbohydrate and collagen) or no feeding control/placebo (5, 7, 8). Furthermore, the European Food Safety Authority (EFSA) advises “human intervention studies assessing the effect of a specific protein source/constituent against another isonitrogenous protein source/constituent were considered as pertinent to the claim, whereas studies controlling for energy only (e.g. using isocaloric carbohydrate sources as placebo) could not be used for the scientific substantiation of these claims” (9) as comparisons of a test protein to isoenergetic, but not isonitrogenous carbohydrate control is more likely to show an effect of protein supplementation (6). Therefore, validation of appropriate non-bioactive isonitrogenous controls is important for the future evaluation of bioactivity of protein formulations.

Studies assessing the role of protein and amino acids in the regulation of MPS establish a close relationship between the extracellular concentration of essential amino acids (EAA) and the rate of MPS (2, 10, 11), and leucine as the most potent in *in vitro* (12) and human studies (8, 13-16). The efficacy of protein and/or amino acid intake to stimulate MPS thereby depends on the pattern and magnitude of change in extracellular EAA evoked following ingestion that, in turn, is dependent on the type, amount and timing of protein or amino acid ingestion. Previous studies also indicate that co-ingestion of non-essential amino acids (NEAA) is surplus to requirement to stimulate MPS (1-4, 17) which is, perhaps, unsurprising as (NEAA) are considered readily available in plasma, interstitial and intracellular muscle compartments. Therefore, a balanced mix of NEAA may also act as an appropriate isonitrogenous control (null) to assess the effect of specific proteins on MPS in humans.

We have recently developed a muscle cell-based model to evaluate the MPS response to ingestion of milk proteins and their derivatives (18). In this model, conditioning media with human serum resulted in an increase in *de novo* MPS in fully differentiated C2C12 skeletal muscle cells. Furthermore, it was also possible to demonstrate that conditioning media with human serum sampled 60 min post-ingestion of whey protein stimulated MPS in mature C2C12 myotubes to a greater extent than serum sampled after an overnight fast (18). Whey is a high EAA (∼50% EAA) containing, soluble milk protein proven to stimulate MPS in young and elderly populations (19, 20). Ingestion of ∼ 0.33g.kg^-1^ body mass whey protein results in post-ingestion aminoacidaemia and an increase in circulating EAA of approximately 3-fold within 60 min (18). Though equivocal, it seemed plausible to suggest the augmented MPS response of myotubes exposed to ‘fed’ *vs*. ‘fasted’ serum conditioned media was due to the increase in circulating EAA in fed serum and that a similar aminoacidaemia through increase in [NEAA] would not result in an increase in MPS. To test this hypothesis, we fed a NEAA formulation designed by Norton et al. (*in review*), isonitrogenous to whey protein, to human participants. Using the C2C12 *in vitro* model established previously (18), serum sampled pre- and post-feeding was used to condition media of C2C12 myotubes to evaluate change in intracellular signalling and *de novo* MPS. The purpose of this study was to evaluate i) if treating skeletal muscle cells with media conditioned by human serum fed a NEAA formulation resulted in increased intracellular signalling and *de novo* MPS and (ii) if media conditioned by human serum fed a NEAA formulation was comparable to the effect of media conditioned by human serum fed WPH.

## 2. Materials and Methods

### 2.1 Ethical Approval

The study was approved by the local ethics committee at the University of Limerick (EHSREC_2013_01_13) and conformed to the standards set by the Declaration of Helsinki. Six young healthy male participants, (26±4.7 y; 77.7±10.1 kg, 1.77±0.08 m, 25±3.3 kg·m^-2^, 19±6.9 % BF) agreed to participate in the study, gave informed written consent and completed the intervention trial.

### 2.2 Study design

Participants reported to the lab following an overnight fast (>10 h) and having not exercised in the previous 24 h on two separate occasions, separated by at least 7 d. A blood sample from the antecubital vein was collected at baseline (t=0 min) by a clinical nurse on each day as described previously (18). Administered double blind participants consumed 0.33 g.kg^-1^ body mass of either an isonitrogenous non-bioactive NEAA control beverage or a WPH (500 mL; 7.6% w/v) beverage within 5 min (Table 1). As aminoacidemia and MPS have been previously shown to peak between 45-90 min following protein feeding (21, 22), an additional blood sample was collected 60 min postprandial.

**Table 1.**
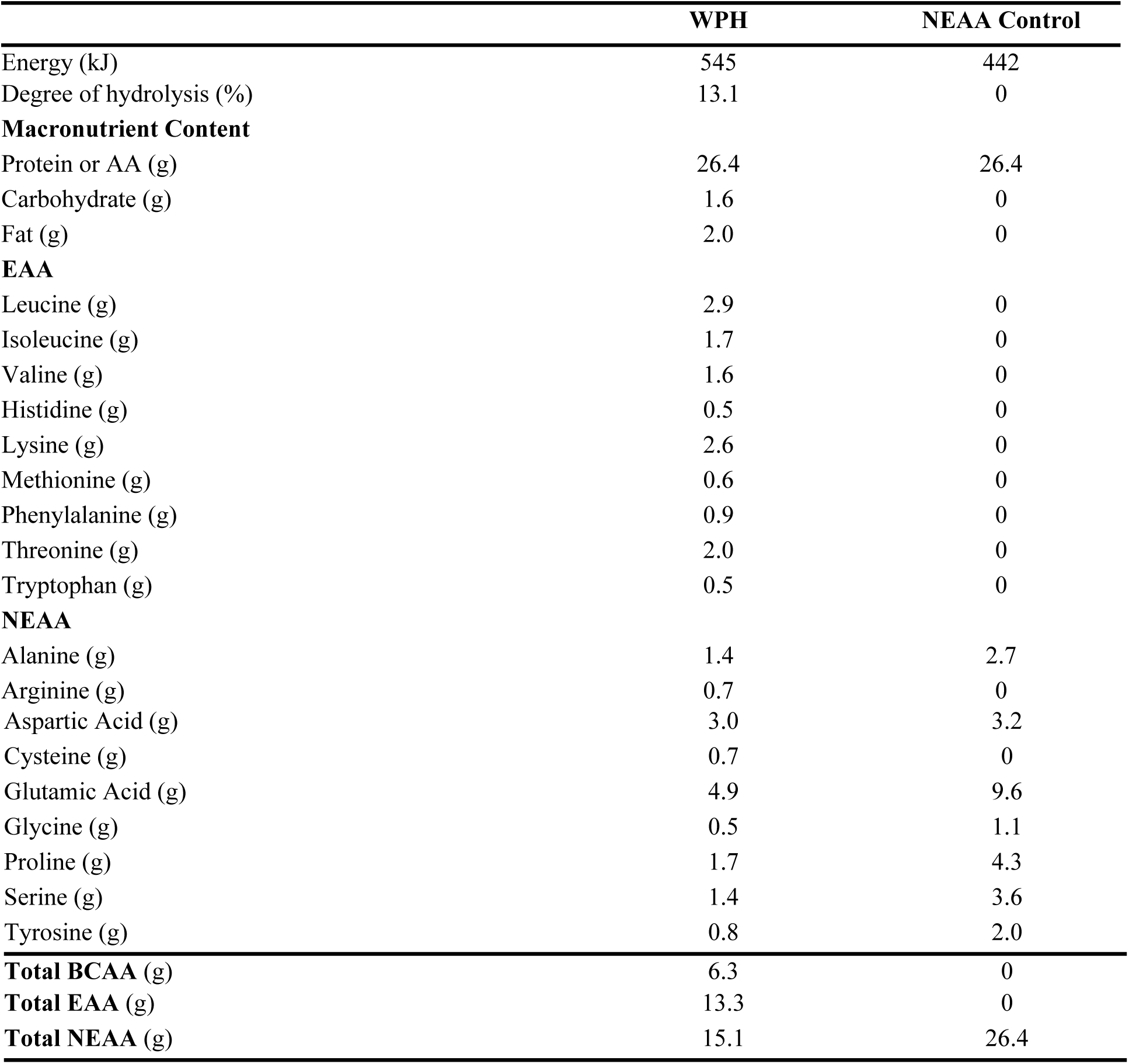
Composition of WPH and isonitrogenous NEAA. Dose scaled per kg body mass (0.33 g.kg^-1^), and amount reported typical for an 80 kg participant. BCAA, branched chain amino acids; EAA, essential amino acids; NEAA, non-essential amino acids; AA, amino acids.

|  | WPH | NEAA Control |
| --- | --- | --- |
| Energy (kJ) | 545 | 442 |
| Degree of hydrolysis (%) | 13.1 | 0 |
| **Macronutrient Content** |  |  |
| Protein or AA (g) | 26.4 | 26.4 |
| Carbohydrate (g) | 1.6 | 0 |
| Fat (g) | 2.0 | 0 |
| **EAA** |  |  |
| Leucine (g) | 2.9 | 0 |
| Isoleucine (g) | 1.7 | 0 |
| Valine (g) | 1.6 | 0 |
| Histidine (g) | 0.5 | 0 |
| Lysine (g) | 2.6 | 0 |
| Methionine (g) | 0.6 | 0 |
| Phenylalanine (g) | 0.9 | 0 |
| Threonine (g) | 2.0 | 0 |
| Tryptophan (g) | 0.5 | 0 |
| **NEAA** |  |  |
| Alanine (g) | 1.4 | 2.7 |
| Arginine (g) | 0.7 | 0 |
| Aspartic Acid (g) | 3.0 | 3.2 |
| Cysteine (g) | 0.7 | 0 |
| Glutamic Acid (g) | 4.9 | 9.6 |
| Glycine (g) | 0.5 | 1.1 |
| Proline (g) | 1.7 | 4.3 |
| Serine (g) | 1.4 | 3.6 |
| Tyrosine (g) | 0.8 | 2.0 |
| **Total BCAA (g)** | **6.3** | **0** |
| **Total EAA (g)** | **13.3** | **0** |
| **Total NEAA (g)** | **15.1** | **26.4** |

### 2.3 Amino acid analysis

Plasma amino acid (AA) profile of each participant at 0 and 60 min postprandial was determined as reported previously (23) on the Agilent 1200 RP-UPLC system (Agilent Technologies, Santa Clara, CA, USA) equipped with an Agilent 1260 binary pump and a G1367C automated liquid handling system. AA separation, data acquisition and quantitative analysis was performed as discussed previously (18).

### 2.4 Metabolic/Humoral biomarker analysis

Plasma insulin was determined using a commercial kit (Merck Millipore) on a MAGPIXTM Multiplex reader and processed using Bio-Plex ManagerTM MP.

### 2.5 Cell culture

*In vitro* analysis was carried out using the murine skeletal muscle cell line C2C12. Cells were cultured, sub-cultured and differentiated (up to 7 d) in DMEM medium as previously described (18, 24). Prior to treatment with media conditioned by human serum, fully differentiated myotubes were nutrient deprived in an AA and serum free DMEM medium (US biological, Salem, MA, USA), supplemented with 1 mM sodium pyruvate (GE Healthcare, Thermo-Fisher), 1% (v/v) penicillin/streptomycin solution, 1 mM L-glutamine, 6 mM D-glucose (Sigma-Aldrich), and 34 mM NaCl (Sigma-Aldrich) (pH 7.3).

### 2.6 Muscle Protein Synthesis

The surface sensing of translation technique (SUnSET) (25) was used to measure muscle protein synthesis in C2C12 myoutbes following treatment with media conditioned by *ex vivo* human serum (fed or fasted). Differentiated and mature C2C12 myotubes were nutrient deprived in AA and serum free DMEM medium for 1 h, following which they were treated with media containing 20% human serum (fed or fasted) and 1 µM puromycin (Merck Millipore Limited) for a further 4 h. The optimum nutrient deprivation time, puromycin, conditioned media treatment time and percentage human serum used has been established previously (18). MPS and protein signalling was determined from cell lysates as described previously (18, 24) and MPS and protein signalling was determined using immunoblotting. Mammalian target of rapamycin complex 1 (mTOR), ribosomal S6 kinase (P70S6K) and eukaryotic initiation factor 4E binding protein 1 (4E-BP1) were the chosen protein signalling targets.

### 2.7 Immunoblotting

Protein lysates (30 μg) were denatured and separated by gel electrophoresis using 4-15% linear gradient SDS-PAGE precast gels (Mini-Protean TGX Stain-free, Bio-Rad 456-8083). Following electrophoresis, total protein (loading control) in each lane of the gel was determined using stain-free UV-induced fluorescence that activates tryptophan residues on the gel (UVITEC Cambridge Imaging system, UVITEC, Cambridge, UK). Semi-dry transfer technique (Trans-blot® Turbo™ Bio-Rad) was adapted to transfer proteins from the UV-activated gel onto a 0.2 μm nitrocellulose membrane. Membranes were probed with the primary antibodies for puromycin (MABE343 anti-puromycin, clone 12D10 mouse monoclonal, Merck Millipore Limited), phosphorylated-4E-BP1 (Thr37/46), 4E-BP1, phosphorylated-P70S6K (Thr389), P70S6K, phosphorylated-mTOR (Ser2448), mTOR and the reference protein β-actin (Cell Signaling). Protein quantification was determined using fluorescence, and membranes were probed with IRDye® 800CW anti-rabbit secondary antibody (926-32211, LI-COR Biosciences UK Ltd, UK) for all proteins except puromycin where IRDye® 800CW goat anti-mouse IgG2a-specific (LI-COR Biosciences UK Ltd) secondary antibody was used. Images were captured in the UVITEC Cambridge Imaging system (UVITEC, Cambridge, UK) and single band and whole-lane (puromycin and total protein) band densitometry was conducted using NineAlliance UVITEC Software (UVITEC, Cambridge, UK).

### 2.8 Statistical analysis

GraphPad Prism v7.03 was used for statistical analysis. Data were tested for normality (Shapiro-Wilk test) and homogeneity of variances (Levene’s test). Paired Samples *T*-tests were used to analyse differences in the plasma [insulin] and [AA] between fasted and fed human condition. Paired sample *T-*Test was used to establish if media conditioned by human serum sampled post-ingestion of the NEAA formulation resulted in increased intracellular signalling and *de novo* MPS in C2C12 myotubes and un-paired sample *T-*Test to establish if these effects were different to the effect of media conditioned by human serum sampled post-ingestion of WPH. The level of significance was set at 95 % (*p* < 0.05).

## 3. Results

Plasma [insulin] and [AA] following an overnight fast and 60 min following WPH or NEAA ingestion (0.33 g.kg^-1^ body mass) are presented in Table 2. Relative to fasting, {AA] increased significantly following WPH ingestion (p<0.05) and only[NEAA] and [threonine]increased following ingestion of NEAA (p<0.05) (**Table 2**). Total [EAA] increased by 97% following WPH ingestion but remained at fasted levels following ingestion of NEAA. Total [NEAA] increased 34% and 55% following WPH and NEAA ingestion, respectively. Plasma [insulin] increased by 44% following WPH (p<0.05) and 41% following NEAA ingestion (p<0.05).

**Table 2.** Plasma insulin and amino acid at baseline (0 min) and postprandial (60 min). EAA, essential amino acids; NEAA, non-essential amino acids.

|  | WPH |  |  | NEAA Control |  |  |
| --- | --- | --- | --- | --- | --- | --- |
| Time (min) | 0 | 60 | Δ (0-60) | 0 | 60 | Δ (0-60) |
| **Humoral Biomarkers** |  |  |  |  |  |  |
| Insulin (pM) | 92 ± 33 | 133±38* | 41 ± 14 | 67 ± 26 | 95 ± 31* | 28 ± 6 |
| **EAA (μmol/L)** |  |  |  |  |  |  |
| Leucine | 127 ± 5 | 333 ± 11* | 207 ± 11 | 151 ± 13 | 138 ± 10 | -13 ± 7<sup>#</sup> |
| Isoleucine | 63 ± 5 | 195 ± 11* | 133 ± 8 | 74 ± 8 | 67 ± 6 | -7 ± 4<sup>#</sup> |
| Valine | 228 ± 7 | 398 ± 10* | 170 ± 9 | 246 ± 16 | 242 ± 12 | -5 ± 9<sup>#</sup> |
| Histidine | 77 ± 2 | 91 ± 4* | 14 ± 3 | 83 ± 5 | 85 ± 4 | 2 ± 3<sup>#</sup> |
| Lysine | 183 ± 8 | 363 ± 17* | 180 ± 13 | 172 ± 14 | 183 ± 14 | 11 ± 6<sup>#</sup> |
| Methionine | 22 ± 3 | 45 ± 2* | 23 ± 4 | 26 ± 2 | 26 ± 2 | 0 ± 1<sup>#</sup> |
| Phenylalanine | 53 ± 2 | 71 ± 3* | 18 ± 1 | 57 ± 5 | 51 ± 11* | -6 ± 2<sup>#</sup> |
| Threonine | 117 ± 12 | 230 ± 14* | 112 ± 5 | 108 ± 9 | 151 ± 11* | 44 ± 4<sup>#</sup> |
| Tryptophan | 67 ± 4 | 124 ± 7* | 57 ± 5 | 72 ± 8 | 71 ± 6 | -1 ± 4<sup>#</sup> |
| **Total EAA (μmol/L)** | 936 ±28 | 1849 ±45* | 914 ± 41 | 989 ± 67 | 1013 ± 55 | 24 ± 37<sup>#</sup> |
| **NEAA (μmol/L)** |  |  |  |  |  |  |
| Alanine | 267 ± 33 | 416 ± 40* | 148 ± 12 | 326 ± 37 | 571 ± 56* | 245 ± 20<sup>#</sup> |
| Arginine | 76 ± 5 | 127 ± 9* | 52 ± 7 | 80 ± 9 | 79 ± 8 | -1 ± 5<sup>#</sup> |
| Asparagine | 125 ± 8 | 187 ± 12* | 62 ± 5 | 131 ± 20 | 255 ± 18* | 125 ± 22<sup>#</sup> |
| Aspartic acid | 0 ± 0 | 4 ± 1* | 3 ± 1 | 0 ± 0 | 11 ± 2* | 11 ± 2<sup>#</sup> |
| Glutamine | 547 ± 20 | 653 ± 37* | 106 ± 20 | 616 ± 33 | 746 ± 43* | 131 ± 23 |
| Glutamic acid | 48 ± 11 | 63 ± 7* | 14 ± 6 | 36 ± 8 | 137 ± 13* | 101 ± 12<sup>#</sup> |
| Glycine | 215 ± 20 | 248 ± 24* | 33 ± 6 | 186 ± 17 | 328 ± 30* | 143 ± 15<sup>#</sup> |
| Tyrosine | 62 ± 4 | 100 ± 6* | 38 ± 3 | 60 ± 7 | 98 ± 11* | 38 ± 5 |
| **Total NEAA (μmol/L)** | 1341 ± 59 | 1798 ± 72* | 457 ± 18 | 1433 ± 92 | 2227 ± 126* | 793 ± 66<sup>#</sup> |
| **Total AA (μmol/L)** | 2276 ± 81 | 3647 ± 67* | 1371 ± 33 | 2422 ± 149 | 3239 ± 175* | 817 ± 97<sup>#</sup> |

*Data are Mean ± SEM (n=6); NA: Not Available; \*within groups $p<0.05$ and <sup>#</sup>between groups $p<0.05$*

Cells treated with media conditioned by WPH-fed serum observed significantly increased mTOR activation (*p*<0.01) (**Figure 1**) relative to its corresponding fasted serum (**Figure 1A, 1B**). This increase in mTOR activation was consistently observed in serum from each participant (n=6) (**Figure 1A**). In comparison, NEAA-fed serum did not activate mTOR (**Figure 1**) which remained at a similar level to its corresponding fasted serum (**Figure 1A, 1B**). Furthermore, when normalised to corresponding fasted serum (**Figure 1B)** mTOR activation in the WPH-fed condition was significantly increased (*p*<0.01) compared to the NEAA-fed condition.

**Figure 1.**
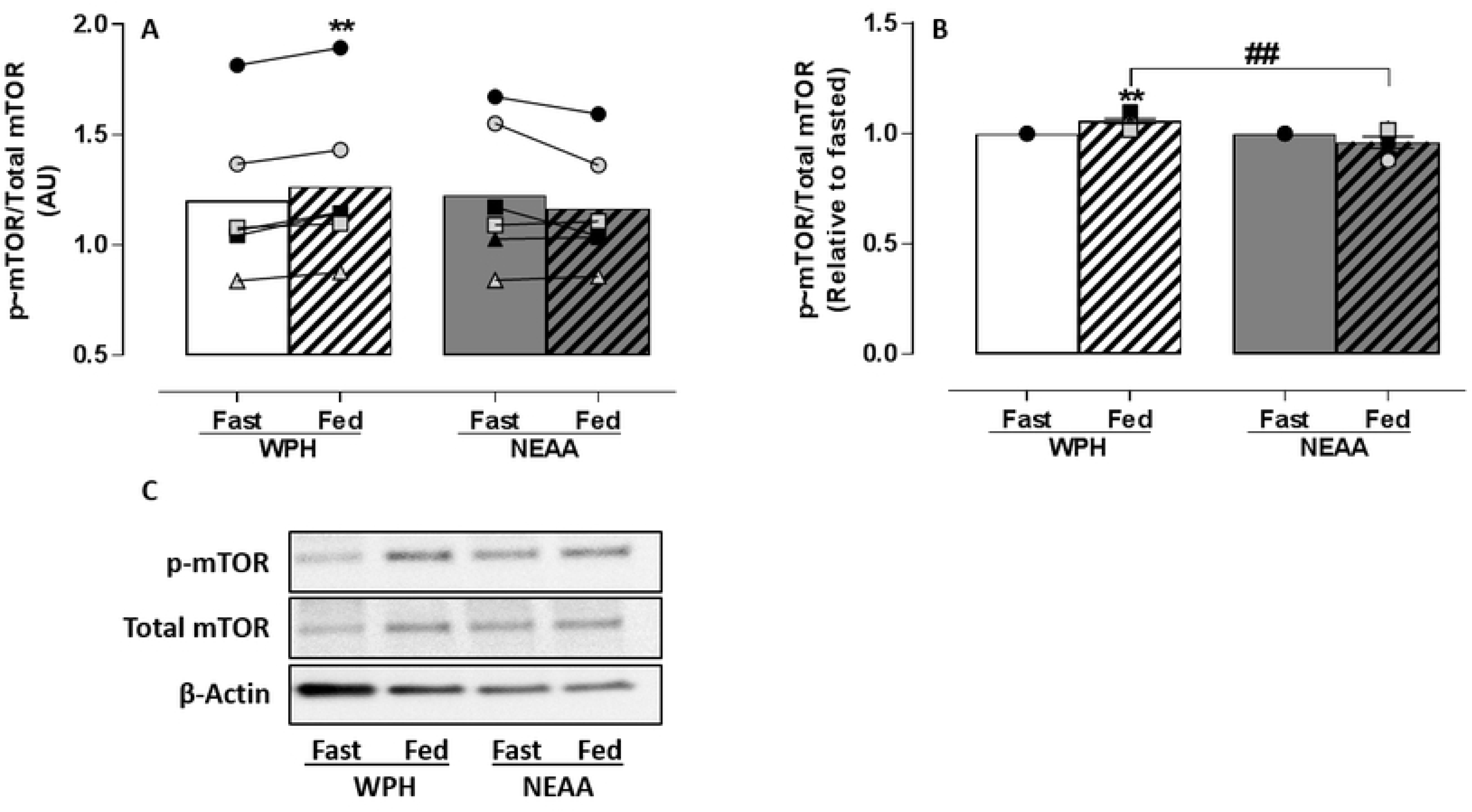
Phosphorylation of mTOR in response to treatment with media conditioned by *ex vivo* human serum (n=6). C2C12 myotubes were nutrient deprived for 1 h followed by treatment with media conditioned by fasted (**fast**) or 60 min postprandial (**fed**) *ex vivo* serum for 4 h. Postprandial serum was obtained 1 h after ingesting WPH or isonitrogenous NEAA. Densitometric analysis of (**A**) mTOR phosphorylation before and after treatment with media conditioned by WPH or NEAA-fed *ex vivo* serum and (**B**) relative to fasted *ex vivo* serum. (**C**) Representative immunoblot of mTOR phosphorylation relative to total mTOR and β-Actin. Data reported as Mean±SEM, \*\**within groups p*<0.01, ^##^*between groups p*<0.01

Stimulation of the downstream targets of mTOR activation, P70S6K and 4E-BP1, occurred following treatment of C2C12 myotubes with media conditioned by serum (**Figure 2**). Like mTOR, P70S6K (**Figure 2A, 2B**) and 4E-BP1 (**Figure 2C, 2D**) activation was significantly increased (*p*<0.05) in the WPH-fed condition relative to its corresponding fasted serum. Comparatively, no change in P70S6K (**Figure 2A, 2B**) or 4E-BP1 (**Figure 2C, 2D**) phosphorylation relative to fasted, following treatment with media conditioned by NEAA-fed serum was evident. This was consistently observed in each participant (**Figure 2A, 2C**). A significant increase (*p*<0.05) in P70S6K (**Figure 2B**) and 4E-BP1 (**Figure 2D**) phosphorylation was observed in media conditioned by WPH-fed compared to NEAA-fed serum (Unpaired T-test).

**Figure 2.**
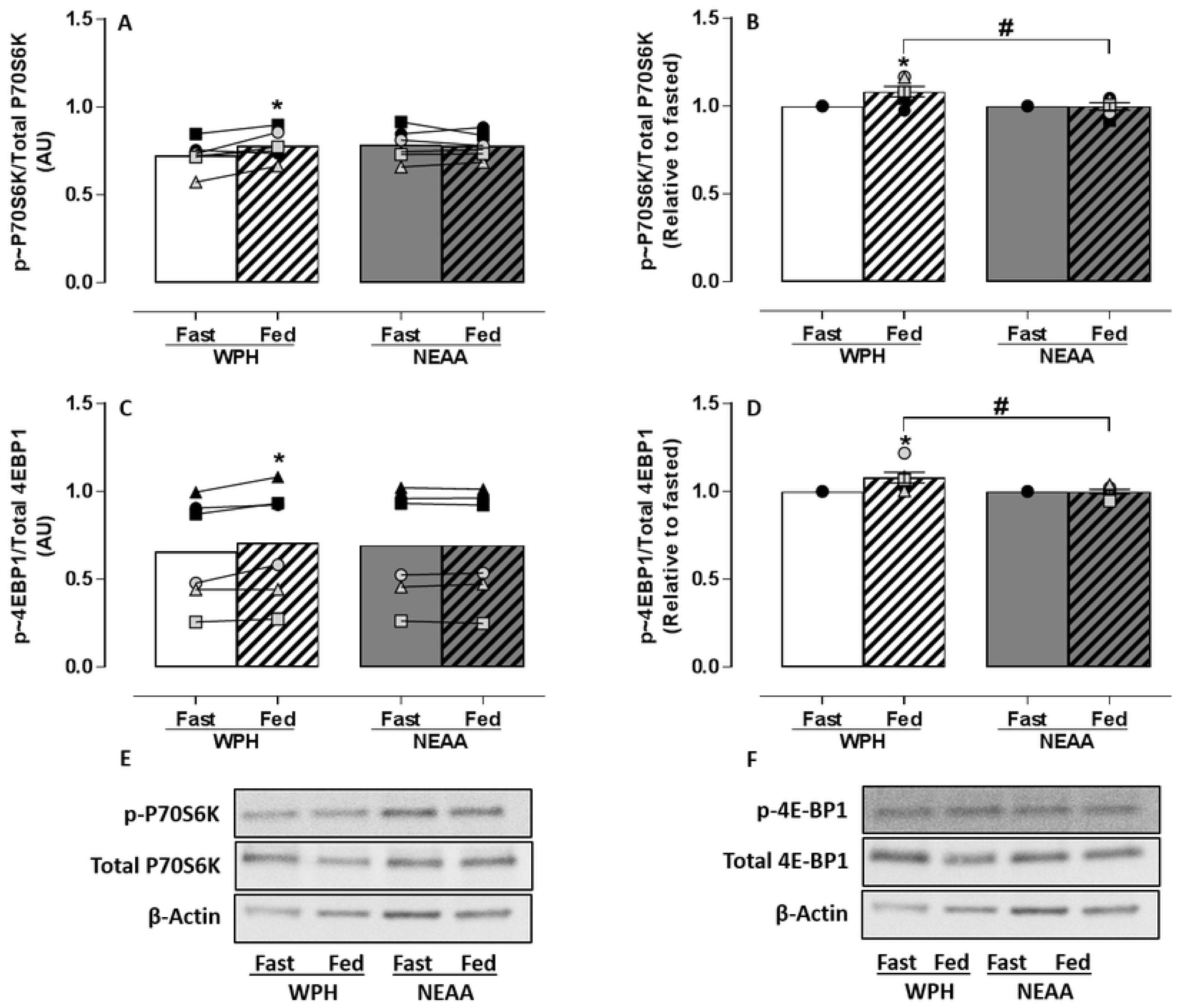
Phosphorylation of P70S6K and 4E-BP1 in response to treatment with media conditioned by *ex vivo* human serum (n=6). C2C12 myotubes were nutrient deprived for 1 h followed by treatment with media conditioned by *ex vivo* fasted (**fast**) or 60 min postprandial (**fed**) serum for 4 h. Postprandial serum was obtained 1 h after ingesting WPH or isonitrogenous NEAA. Densitometric analysis of P70S6K and 4E-BP1 phosphorylation before and after treatment with media conditioned by WPH or NEAA-fed serum (**A, C**) and relative to fasted serum (**B, D**). Representative immunoblots of P70S6K (**E**) and 4E-BP1 (**F**) phosphorylation relative to their respective total proteins and β-Actin. Data reported as Mean±SEM, \**within groups p*<0.05, ^#^*between groups p*<0.05

The SunSET technique (25) was adopted to verify that mTOR, P70S6K and 4E-BP1 activation led to an increase in *de novo* MPS in skeletal muscle cells. No statistically significant increase in MPS occurred in skeletal muscle cells treated with media conditioned by NEAA-fed or WPH-fed serum when compared to treatment with media conditioned by their corresponding fasted serum (**Figure 3**). However, normalised to the corresponding fasted serum, significantly greater MPS was detected in the cells treated with media conditioned by WPH-fed compared with NEAA-fed serum (*p*<0.05) (**Figure 3B**).

**Figure 3.**
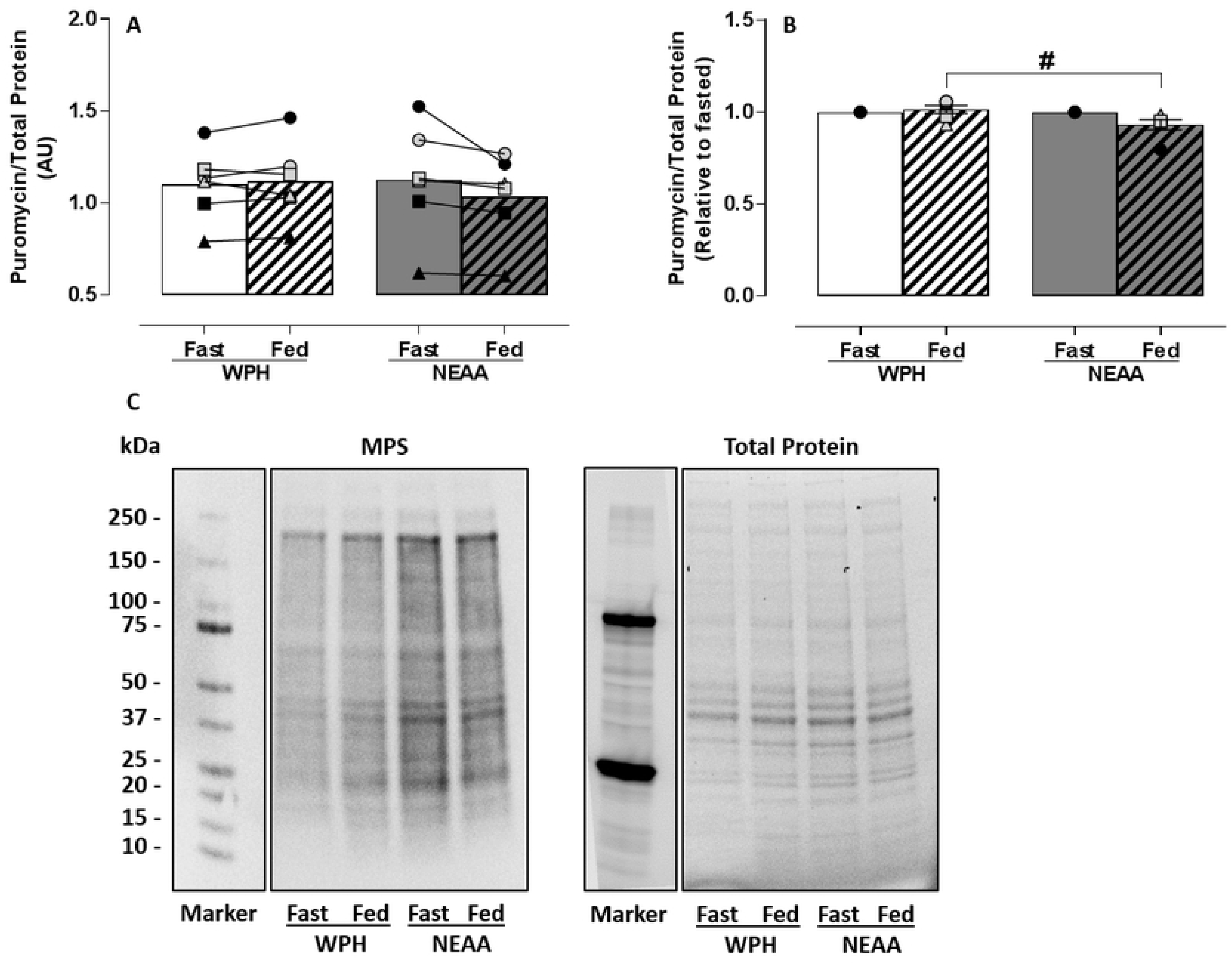
MPS in response to treatment with media conditioned by *ex vivo* human serum (n=6). C2C12 myotubes were nutrient deprived for 1 h followed by treatment with *ex vivo* fasted (**fast**) or 60 min postprandial (**fed**) human serum for 4 h. Postprandial serum was obtained 1 h after ingesting WPH or isonitrogenous NEAA. Densitometric analysis of (**A**) MPS before and after treatment with media conditioned by WPH or NEAA-fed serum and (**B**) relative to fasted serum. (**C**) Representative immunoblot of MPS (measured by puromycin incorporation) relative to total protein (loading control). Data reported as Mean±SEM, ^#^*between groups p*<0.05

## 4. Discussion

The importance of the use of appropriate negative controls in human MPS studies has recently been emphasised by key opinion leaders in the field (5, 6). Many studies in these meta-analyses report on the effects of amino acid and protein supplementation on MPS in humans which use less appropriate controls (5, 7, 8). In line with the scientific opinion of EFSA, that human intervention studies assessing the effect of different proteins on physiological processes require an isonitrogenous comparator, a NEAA-only containing formulation isonitrogenous to whey protein, was fed to human participants in equal dose to a whey protein hydrolysate. Fed and fasted serum was used to condition media of C2C12 myotubes and evaluate change in intracellular signalling and *de novo* MPS. Our findings show that media conditioned by WPH-fed serum stimulated kinases of the mTOR pathway and MPS *in vitro*, however media conditioned by NEAA-fed serum did not.

As expected, plasma EAA concentration, including leucine, were not significantly elevated from fasting levels following NEAA ingestion, but were increased following WPH ingestion (**Table 2**). Elevated plasma levels of EAA post protein feeding have previously been shown to robustly increase MPS (15, 26). Rennie and colleagues reported that an increase of ∼80 µmol/L in extracellular leucine is required to increase MPS *in vivo* (27, 28). Here, we observed an increase greater than ∼200µmol/L for WPH-fed, however, this threshold was not reached in NEAA-fed condition and likely explains the lack of activation of MPS in our model, confirming NEAA as an effective isonitrogenous non-bioactive control.

Several humoral factors may act individually or collectively to stimulate MPS. Whereas circulating EAA are thought to be the primary drivers of MPS, protein ingestion has been shown to induce an increase in insulin, which is deemed ‘permissive’ with respect to MPS (29). In this study, we report a small increase (∼40 - 45%) in circulating insulin with ingestion of both WPH and NEAA. As reviewed elsewhere (30) and following a recent meta-analysis (31), large increases in MPS are due to EAA regulating anabolic responses, whereas insulin regulates anti-catabolic (MPB) responses independent of AA availability (30, 31). Insulin, even in low concentrations as observed here (WPH: 133 ± 38; NEAA 95 ± 31), has been shown to attenuate MPB *in vivo* (32), however, this is not thought to impact MPS (the focus of this paper) when EAA delivery is not increased as in the case of the non-bioactive NEAA control. Consumption of NEAA did result in insulin-mediated clearance of EAAs from the circulation with small reductions in circulating EAAs ranging from 1-10 %. As a result, we postulate that only an elevation in circulating EAA would result in increased signalling and MPS in response to media conditioned by *ex vivo* protein-fed serum. Therefore, in the absence of an increase in circulating EAA as observed here, we anticipate that an isonitrogenous NEAA formulation can act as an effective non-bioactive control for further investigation of the effect of protein feeding on MPS in this model.

The potential of media conditioned by *ex vivo* human serum fed with isonitrogenous NEAA or WPH to activate MPS in C2C12 myotubes was measured by phosphorylation of mTOR and its downstream molecular proteins P70S6K and 4E-BP1. Dickinson and colleagues determined that activation of mTOR and its downstream signalling proteins P70S6K and 4E-BP1 is required for stimulation of human skeletal MPS by EAA (33). Addition of NEAA-fed serum to condition cell media did not stimulate mTOR, P70S6K or 4E-BP1 phosphorylation in C2C12 skeletal muscle cells. This confirms a lack of bioactivity for the activation of MPS in the NEAA formulation. In comparison, media conditioned by WPH-fed serum significantly increased phosphorylation of mTOR, P70S6K and 4E-BP1, expressed as absolute values (**Figure 1A, 2A, 2C**), normalised relative to fasted serum (**Figure 1B, 2B, 2D**) and in comparison to NEAA-fed. These *in vitro* data confirm the bioactivity of WPH to activate MPS. Furthermore, activation of mTOR, P70S6K and 4E-BP1 with media conditioned by WPH-fed serum resulted in significantly greater stimulation of *de novo* MPS than media conditioned by NEAA-fed serum (**Figure 3B**), providing further validation of the NEAA formulation used in this study as an isonitrogenous, non-bioactive control.

In this study, we have demonstrated that an isonitrogenous NEAA supplement can be used as a non-bioactive control for MPS in protein feeding studies. As discussed, NEAA have previously been demonstrated not to be primarily responsible or required to stimulate MPS (1-3, 17). Similarly, unlike a bioactive WPH supplement, the isonitrogenous non-bioactive NEAA control used here did not alter protein signalling activity of mTOR, P70S6K and 4E-BP1 or MPS levels relative to its corresponding fasted serum. This suggests that in acute feeding studies, this isonitrogenous non-bioactive NEAA supplement can serve as an appropriate control.

## 5. Conclusions

In conclusion, we have proposed and demonstrated the use of an isonitrogenous NEAA control that does not affect levels of circulating biomarkers, does not mediate signalling through the mTOR pathway and neither augments nor attenuates MPS when used to condition media of skeletal muscle cells *in vitro*. We have also demonstrated that an isonitrogenous non-bioactive NEAA control can be used as a comparator against a bioactive WPH in this model. This study also provides further evidence on the use of pre- and post-fed *ex vivo* human serum in regulating MPS *in vitro*.

## 6. Acknowledgements

This work was supported by Food for Health Ireland (Enterprise Ireland grant TC20130001 to PMJ & BPC).

## Author Contributions

BPC, PMJ: conception, design, interpretation of data, drafting and revising the manuscript critically for important intellectual content. RJF, ABN: BP, MP, MAB: data acquisition, analysis, interpretation of data, drafting and revising of the manuscript.

